# Amino acid quality modifies the quantitative availability of protein for reproduction in *Drosophila melanogaster*

**DOI:** 10.1101/2020.03.26.010538

**Authors:** Carolyn Ma, Christen K. Mirth, Matthew D. Hall, Matthew D.W. Piper

## Abstract

Diet composition, especially the relative abundance of key macronutrients, is well known to affect animal wellbeing by changing reproductive output, metabolism and length of life. However, less attention has been paid to the ways the quality of these nutrients modify these macronutrient interactions. Nutritional Geometry can be used to model the effects of multiple dietary components on life-history traits and to compare these responses when diet quality is varied. Previous studies have shown that dietary protein quality can be increased for egg production in *Drosophila melanogaster* by matching the dietary amino acid proportions to the balance of amino acids used by the sum of proteins in the fly’s *in silico* translated exome. Here, we show that dietary protein quality dramatically alters the effect of protein quantity on female reproduction across a broad range of diets varying in both protein and carbohydrate concentrations. These data show that when sources of ingredients vary, their relative value to the consumer can vastly differ and yield very different physiological outcomes. Such variations could be particularly important for meta analyses that look to draw generalisable conclusions from diverse studies.

## Introduction

To optimise fitness, organisms must consume a sufficient quantity and quality of nutrients to suit their needs (Hall et al., 2008, Lee et al., 2008, Bong et al., 2014). This is demonstrated by the dramatic changes in reproductive function seen when flies or rodents are subjected to food restriction or changes in diet balance, particularly when the relative proportion of protein to carbohydrates is changed (Widdowson and Cowen, 1972, Good and Tatar, 2001, Carey et al., 2002, Liang and Zhang, 2006, Lee et al., 2008, Skorupa et al., 2008, Simpson and Raubenheimer, 2012, Solon-Biet et al., 2015, Camus et al., 2019).

Food is comprised of dozens of nutrients that interact to modify animal physiology. Understanding how nutritional interactions affect the consumer is complex but can be facilitated using a structured approach such as Nutritional Geometry (Raubenheimer and Simpson, 1997). Nutritional Geometry maps the responses of life history traits to quantitative variations of two or more macronutrients. This is typically performed by exposing animals to one of many diets that vary across a range of calorie compositions and protein to carbohydrate ratios. A nutrient space is defined by axes (generally two) that represent the quantity of nutrients that an organism has eaten – thus any point in space can represent the status of an organism according to its nutritional history. By mapping an array of organisms with different dietary histories into this space, their collective phenotypic responses can be fitted by an overlaid surface represented by a heat map (*z*-axis). This allows for the modelling of the interactive effects of nutrients on phenotypes of interest. Although conceptually simple, assessing phenotypic responses to nutrition through the perspective of Nutritional Geometry has revealed new understanding of biology and, in some cases, has unified apparently conflicting interpretations about the way organisms respond to diet change (Solon-Biet et al., 2016).

Nutritional Geometry experiments have shown that variation in two of the energy-yielding macronutrients, protein and carbohydrate, affect the expression of many traits. In particular the lifespan and reproduction of adult fruit flies (*Drosophila melanogaster*) are shaped by the interactive effects of dietary protein and carbohydrate (Lee et al., 2008, Skorupa et al., 2008, Jensen et al., 2015). In the case of protein, more recent work has shown that the proportion of its constituent amino acids has an important role to play in protein’s effects on these traits (Grandison et al., 2009, Piper et al., 2017).

The dietary amino acid requirements of female flies for optimal egg production can be determined from its genome by a process termed exome matching (Piper et al., 2017). To exome match a diet, we *in silico* translate the exome of the consumer, sum the abundance of each amino acid across all proteins, and find the relative proportion of each amino acid. We then use this proportion as the basis for the abundance of each amino acid in the food. By matching the dietary protein quality to the fly exome in this way, we found that for a fixed mass of amino acids, exome matched diets supported higher levels of reproduction than diets that were mismatched (Piper et al., 2017). Together, these data show that dietary amino acid balance is important for determining fitness outcomes.

Although dietary amino acid balance is important, Nutritional Geometry experiments that measure fitness responses to diets invariably treat protein as a single nutrient dimension with a fixed proportion of all 20 amino acids. This is reasonable for experiments in which the protein source is held constant across all diets. However, proteins vary in quality when attained from different sources and, like many natural ingredients, the same type of protein may vary in quality between locations and seasons. Given this, it may be difficult to directly compare the effects of consumed “protein” on a trait when the data are from different studies. To examine these effects, we designed an experiment using a Nutritional Geometry design to assess how changing dietary protein quality modifies the interactive effects of the amino acid (protein; P) to carbohydrate (C) ratio on egg laying. We designed an array of diets varying in P:C ratio for each of two different amino acid (protein) mixtures. These mixtures varied in the relative proportion of each amino acid, as well as the identity of the most limiting essential amino acid and the degree to which it is predicted to be limiting.

## Method

### Animal husbandry

We used the Dahomey outbred population of *Drosophila melanogaster* (Mair et al., 2005). Routine rearing and maintenance of flies employed the techniques and sugar/yeast (SY) diet described in Bass et al. (2007). All flies were maintained at 25°C with a 12 hr: 12 hr light dark photoperiod. For the experiment, a population of flies was age synchronised as in Piper and Partridge (2016).

### Experimental diets

Completely defined synthetic (holidic) diets were made according to Piper et al. (2014), in which free amino acids are used to make up protein equivalents. To convert amino acids to protein equivalents, we used the molar quantities of nitrogen and the assumption that N makes up 16% of whole proteins (Sosulski and Imafidon, 1990). Two protein qualities, defined by their amino acid ratios, were compared: FLYaa (matched to the amino acid ratio of the exome of adult flies), and MMaa (a ratio considered mis-matched to the flies’ requirements). The relative proportion of each amino acid within each amino acid ratios are displayed in Table 1. From this point on, the concentration of total amino acids is referred to as protein (P), calculated according to the above method.

**Table 1.** The relative proportions of each amino acid in the two amino acid ratios tested, FLYaa and MMaa.

| Amino acids |  |  | Ratio |  |
| --- | --- | --- | --- | --- |
|  |  |  | MMaa | FLYaa |
| Essential amino acids | Phenylalanine | F | 0.027 | 0.037 |
|  | Histidine | H | 0.022 | 0.026 |
|  | Isoleucine | I | 0.077 | 0.052 |
|  | Lysine | K | 0.044 | 0.057 |
|  | Leucine | L | 0.052 | 0.094 |
|  | Methionine | M | 0.018 | 0.025 |
|  | Arginine | R | 0.016 | 0.057 |
|  | Threonine | T | 0.057 | 0.056 |
|  | Valine | V | 0.081 | 0.062 |
|  | Tryptophan | W | 0.008 | 0.010 |
| Non-essential amino acids | Alanine | A | 0.133 | 0.075 |
|  | Cysteine | C | 0.001 | 0.017 |
|  | Aspartate | D | 0.043 | 0.053 |
|  | Glutamate | E | 0.058 | 0.063 |
|  | Glycine | G | 0.144 | 0.062 |
|  | Asparagine | N | 0.044 | 0.047 |
|  | Proline | P | 0.044 | 0.052 |
|  | Glutamine | Q | 0.058 | 0.046 |
|  | Serine | S | 0.061 | 0.079 |
|  | Tyrosine | Y | 0.013 | 0.031 |

For both amino acid ratios, we generated diets that were one of five P:C ratios (1:3.6, 1:1.8, 1:1.1, 1:0.8, 1:0.6) and one of four caloric densities (66.8 kcal/L, 111.3 kcal/L, 155.8 kcal/L, 200.3 kcal/L), where dietary energy densities were estimated by calculation, using a value of 4 kcal/g for both protein and carbohydrates (Table 2). Thus, 20 diets were employed to test the effect on egg laying of each amino acid ratio.

**Table 2.** The equivalent protein: carbohydrate (P:C) ratio, displayed with the nutrient densities.

| P:C equivalent | Sum mass of amino acids (g/L) | Equivalent protein (g/L) | Equivalent carbohydrate (g/L)<sup>1</sup> | Estimated caloric content (kcal/L) |
| --- | --- | --- | --- | --- |
| 1:3.6 | 3.92 | 3.54 | 12.77 | 66.8 |
| 1:3.6 | 6.53 | 5.91 | 21.29 | 111.3 |
| 1:3.6 | 9.14 | 8.27 | 29.81 | 155.8 |
| 1:3.6 | 11.75 | 10.63 | 38.32 | 200.3 |
| 1:1.8 | 6.42 | 5.81 | 10.27 | 66.8 |
| 1:1.8 | 10.7 | 9.68 | 17.12 | 111.3 |
| 1:1.8 | 14.98 | 13.55 | 23.97 | 155.8 |
| 1:1.8 | 19.26 | 17.42 | 30.82 | 200.3 |
| 1:1.1 | 8.32 | 7.52 | 8.38 | 66.8 |
| 1:1.1 | 13.86 | 12.54 | 13.96 | 111.3 |
| 1:1.1 | 19.41 | 17.55 | 19.54 | 155.8 |
| 1:1.1 | 24.95 | 22.57 | 25.13 | 200.3 |
| 1:0.8 | 9.57 | 8.66 | 7.12 | 66.8 |
| 1:0.8 | 15.95 | 14.43 | 11.87 | 111.3 |
| 1:0.8 | 22.33 | 20.20 | 16.62 | 155.8 |
| 1:0.8 | 28.71 | 25.97 | 21.37 | 200.3 |
| 1:0.6 | 10.99 | 9.94 | 5.70 | 66.8 |
| 1:0.6 | 18.32 | 16.57 | 9.50 | 111.3 |
| 1:0.6 | 25.64 | 23.20 | 13.30 | 155.8 |
| 1:0.6 | 32.97 | 29.65 | 17.10 | 200.3 |

**Table 2.** Egg production on each amino acid ratio (FLYaa and MMaa) was modelled using linear predictors for protein (P) and carbohydrate (C) and their interaction (P x C). Because the data appear non-linear, we also assessed the quadratic terms of protein (P^2^) and carbohydrates (C^2^). Minimum adequate models are reported. *β* indicates the the slope estimate for each variable, and; SE, the standard error. Highlighted bold signifies significance, * *p*<0.05; ** *p*<0.001.

| Amino acid ratio | | P | C | $P^2$ | $C^2$ | P x C |
| --- | --- | --- | --- | --- | --- | --- |
| FLYaa | $\beta$ | **3.250** | -0.206 | **-0.127** | -0.007 | 0.042 |
|  | SE | **1.393** | 1.203 | **0.037** | 0.024 | 0.050 |
|  | t-value | **2.332\*** | -0.171 | **-3.464\*\*** | -0.301 | 0.846 |
| MMaa | $\beta$ | **1.186** | 0.129 | N/A<sup>2</sup> | N/A<sup>2</sup> | -0.032 |
|  | SE | **0.502** | 0.341 |  |  | 0.027 |
|  | t-value | **2.365\*** | 0.377 |  |  | -1.184 |

<sup>2 2</sup> N/A, not applicable. For the MMaa model, $P^2$ and $C^2$ were not significant in the model and so were removed during model reduction.

### Experimental set-up

Flies were housed in devices called dFlats, which are made up of a block of Perspex with 12-wells drilled into them, such that the drilled wells have a volume the same as standard fly vials (FS32, Pathtech) and their arrangement is such that the 12 openings match the position of wells in a 12-well plate (based on the design: https://www.flidea.tech/proiects). 3ml food was dispensed into each of the wells in a 12-well plate. Each dFlat well contained 10 mated female flies and we maintained one dFlat for each of the 20 holidic experimental diets (12 replicate wells x 10 flies = 120 flies per diet).

### Reproduction experiment

Once a week for 3 weeks the number of eggs laid on the media over an 18-hour period was counted and recorded. Measuring reproductive output during the first weeks of egg laying has shown to be representative of life-long reproduction of flies (Chapman and Partridge, 1996, Muller et al., 2001). For each well in each dFlat device, the number of eggs laid per female on each experimental day (day 8, 15 and 22) was calculated. The value for eggs laid by an average female in a well was summed across days and used to calculate the cumulative egg laying of an average female in a food type. We call this value the index of reproduction. The number of eggs in each well and food type was obtained by taking images using a web camera attached to a stereomicroscope. The images were then processed in Image J to acquire the correct image size, which were then automatically counted in the software called QuantiFly (Waithe et al., 2015).

### Statistical analyses

All analyses were conducted using R (version 3.3.0, available from http://www.R-project.org/). To analyse the relationship between our index of reproduction and the protein (P) and carbohydrate (C) concentration, we generated separate response surfaces for each of the two amino acid ratios. Each surface was estimated using multivariate second-order polynomial regression, whereby the linear, quadratic and cross-product terms from this model capture the linear and non-linear effects of P and C concentration on fly reproduction. For each amino acid ratio, the minimum adequate model for each linear model was found by determining if eliminating the most complex parameter significantly reduced the model fit. We visualised the response surface of each amino acid ratio using predictions derived from thin-plate splines from the fields package (see www.github.com/NCAR/Fields) and subsequently visualised via the ggplot package.

## Results

To capture the effects of differences in protein quality on egg production of *D. melanogaster, we* made media containing two different amino acid ratios, one known to be mismatched to fly requirements for egg laying (MMaa) and the other known to be matched precisely to the fly’s exome (FLYaa) (Piper et al., 2017). For each amino acid ratio, an array of 5 different P:C ratios, each at 4 different nutrient densities, was tested. Thin-plate splines were used to visualise egg laying response surfaces of flies maintained on the different diets (Figure 1A and 1B).

**Figure 1.**
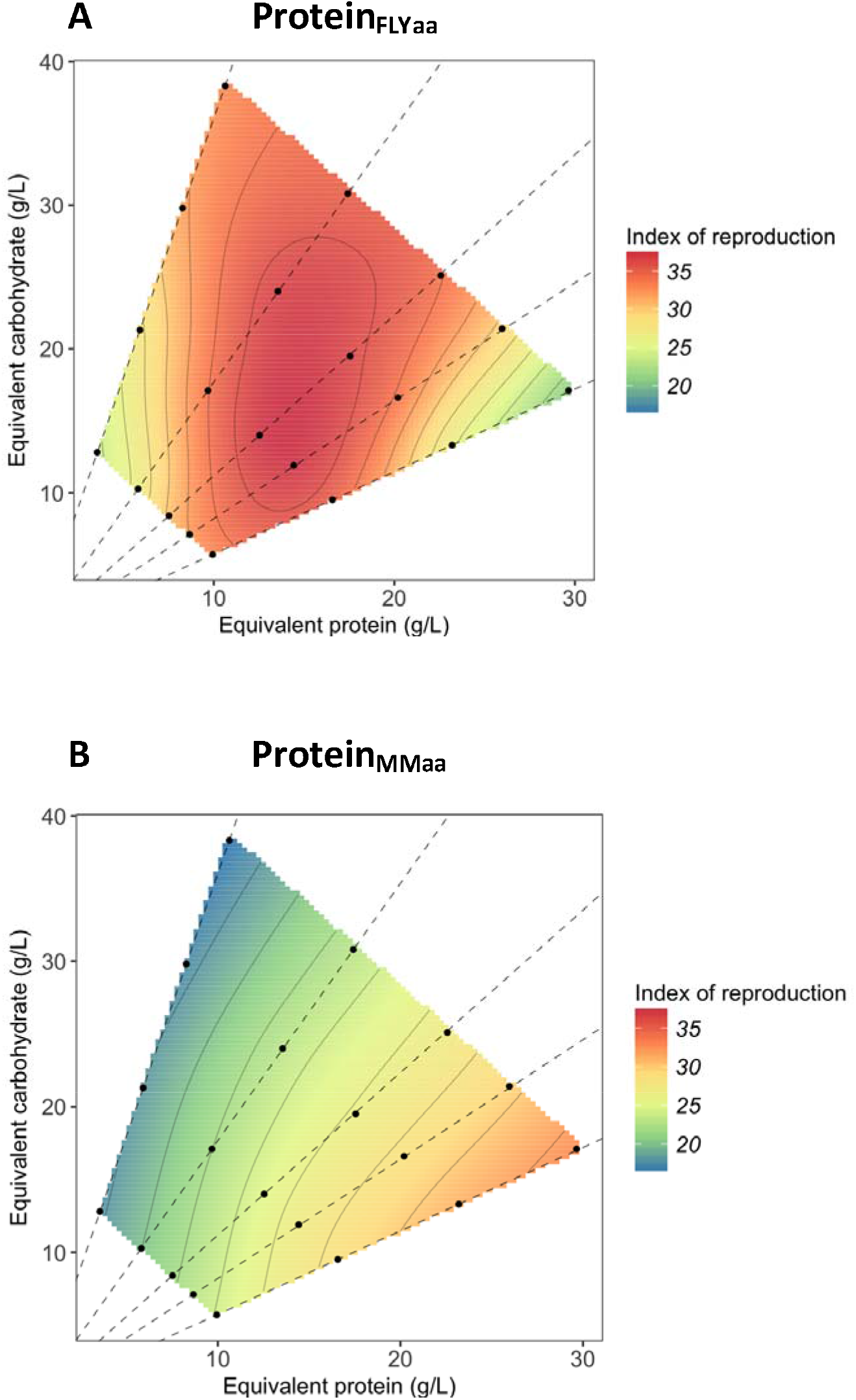
Egg laying varies with the quantity of carbohydrates and the quantity and quality of protein. Flies were maintained on diets containing amino acids in the ratio of either (A) FLYaa (fly exome matched amino acid ratio) or (B) MMaa (mis-matched amino acid ratio). The black dots represent the individual 20 diets from which the index of reproduction was assessed and the dashed lines represent nutritional rails of fixed P:C.

Across all food types, flies maintained on the MMaa diets laid fewer eggs per female per day than those on the FLYaa diets (MMaa diet, 24.07 ± 1.42; FLYaa 31.69 ± 2.14; one-way ANOVA: F_1,33_ = 11.205, p < 0.001). For flies on both the MMaa and the FLYaa diet, there was a significant effect of the linear component of protein, such that egg laying increased with increasing dietary protein concentration (Table 2.). For FLYaa, we also found the quadratic term for protein concentration to be significant, which is shown in the plots as the peak of egg laying in FLYaa occurred at intermediate concentrations of protein (P:C of 1:0.8) (Figure 1a; Table 2). From this point, egg laying dropped away as the protein concentration either increased or decreased (Figure 1a; Table 2). In contrast, peak egg laying on MMaa occurred on the food with the highest nutrient density with P:C of 1:0.6. There was no detectable effect of carbohydrate concentration on egg production across foods of both amino acid ratio and there was also no significant interaction between carbohydrate and protein concentration detected (Table 2). These results show that for the range of diets we tested, protein was the principle determinant of egg production, and that per gram of amino acids supplied, FLYaa supported higher levels of egg laying than MMaa.

## Discussion

The ratio of dietary protein to carbohydrate in the diet has an important role in determining reproduction in *Drosophila melanogaster*: higher density diets with greater P:C ratios support higher female egg laying (Mair et al., 2005, Lee et al., 2008, Skorupa et al., 2008, Jensen et al., 2015). Here we show that varying the protein quality of a diet can also dramatically alter its bioavailability for reproduction of female *D. melanogaster*. These data demonstrate how diverse outcomes in important fitness traits can occur when the quality of food ingredients vary.

Protein is often a limiting component of the diet for terrestrial animals (White, 1993), which means that changes in the quality of dietary protein consumed should have observable effects on fitness for many animals in diverse settings. Indeed, supplementing the diet of wild cotton rats and cottontail rabbits with an essential amino acid (methionine) can improve their reproductive success and increase population size (Lochmiller et al., 1995, Webb et al., 2005), while maintaining blue tits on a diet supplemented with amino acids matched for egg protein formation laid more eggs per clutch than those that received a mismatched balance of amino acids (Ramsay and Houston, 1998). Similarly, female copepods fed on a diet containing essential amino acid profiles that were closely matched to female body composition had higher reproductive success (Guisande et al., 1999) and, supplementing amino acids in the diet of livestock, like boars and chickens, can increase reproductive output and yield more lean muscle (Dong et al., 2016, Cerrate et al., 2019). Finally, recent work in the lab on flies and mice has shown that differences in protein quality, specified at the level of essential amino acid balance, can have dramatic effects on animal reproduction and feeding behaviour (Leitao-Goncalves et al., 2017, Piper et al., 2017, Solon-Biet et al., 2019). Thus, changing the quality of dietary protein by altering the proportion of amino acids can have important effects on fitness traits of both laboratory-reared and wild animals.

In our experimental diets, we found large changes in egg output because the amino acid ratio was altered so that the bioavailability of the nitrogen source varied 2.5-fold (Piper et al., 2017). In other words, the level of FLYaa that was required to support maximal egg output was ∼15g/L, which is 2.5-times less than the amount of MMaa (37 g/L) that would be required to achieve the same output. Thus, smaller amounts of high-quality food are required for optimal egg laying. With defined diets, this large difference in protein quality is easily generated because each amino acid can be independently manipulated over a wide range of concentrations. Interestingly, similarly large differences in the relative abundance of amino acids (i.e. changes in protein quality) can also be found between natural proteins in whole foods. For example, when comparing the average amino acid profiles across food groups published in an FAO report (Food Policy and Food Science Service, 1970), we found that for a fixed quantity of protein, the average amino acid proportions in meat represent ∼2-fold higher quality than the average legume-based protein. This is because the limiting essential amino acid in legume protein, methionine, is more deficient compared to the limiting essential amino acid, leucine,in meat protein. Thus, we predict that using different dietary components to feed animals or humans will result in dramatic changes to physiology and behaviour. Furthermore, without explicit knowledge of the source of ingredients and their quality, it will be difficult to extract generalisable conclusions from meta-analyses that incorporate diverse studies.

It is important to note that protein is not the only compound axis in most published Nutritional Geometry experiments. For example, the other energy yielding macronutrients, carbohydrate and fat, also exist as a diverse array of molecules in food. This can be even more complicated since in typical studies with flies, sucrose and yeast are often used the sources of carbon and protein, respectively (Bass et al., 2007, Min et al., 2007, Lee et al., 2008), and yeast contains various types of carbohydrates as well as other nutritional components, such as sterols, nucleic acids, vitamins, and minerals (Lange and Heijnen, 2001). In addition to our findings about protein quality, other work has shown that varying the identity of the carbohydrate component of the diet from sucrose to fructose can alter fly physiology (Lushchak et al., 2014). Thus, it will be important in future studies to understand the relative contribution of different carbohydrates, fats and other nutrients in modifying fitness traits.

## Conclusion

By showing that the precise amino acid composition of dietary protein is key for dictating female fecundity, we demonstrate the need for more information to be provided when labelling composite nutritional axes in Nutritional Geometry experiments. In particular, protein should be labelled with its source to indicate its quality – a metric that would be further improved if the most limiting essential amino acid were identified by referencing the dietary protein amino acid profile to the *in silico* translated exome of the consumer (Piper et al., 2017). We anticipate this will facilitate the comparability of data between studies.

## Acknowledgements

We thank Amy Dedman, Sasha Pollock and Sarah Gough for technical assistance and Scott Pletcher for help with design and development of the hotel chambers. We also acknowledge many fruitful discussions at the Nutritional Homeostasis workshops in Bonn.

